# Radixin modulates stereocilia function and contributes to cochlear amplification

**DOI:** 10.1101/2020.02.11.944355

**Authors:** Sonal Prasad, Barbara Vona, Marta Diñeiro, María Costales, Rocío González-Aguado, Ana Fontalba, Clara Diego-Pérez, Asli Subasioglu, Guney Bademci, Mustafa Tekin, Rubén Cabanillas, Juan Cadiñanos, Anders Fridberger

**Affiliations:** Department of Biomedicine and Clinical Sciences, Linköping University, SE-581 83 Linköping, Sweden; Department of Otorhinolaryngology, Head and Neck Surgery, Tübingen Hearing Research Centre, Eberhard Karls University Tübingen, 72076 Tübingen, Germany; Laboratorio de Medicina Molecular, Instituto de Medicina Oncologica y Molecular de Asturias, 33193 Oviedo, Spain; Department of Otorhinolaryngology, Hospital Universitario Central de Asturias, 33011 Oviedo, Spain; Department of Otorhinolaryngology, Hospital Universitario Marqués de Valdecilla, 39008 Santander, Spain; Department of Genetics, Hospital Universitario Marqués de Valdecilla, 39008 Santander, Spain; Department of Otorhinolaryngology, Hospital Universitario de Salamanca, 33007 Salamanca, Spain; Department of Medical Genetics, Izmir Ataturk Education and Research Hospital, Izmir 35360, Turkey; John P. Hussman Institute for Human Genomics, University of Miami Miller School of Medicine, Miami, FL 33136, USA; Department of Otolaryngology, University of Miami Miller School of Medicine, Miami, FL 33136, USA; Dr. John T. Macdonald Department of Human Genetics, University of Miami Miller School of Medicine, Miami, FL 33136, USA; Área de Medicina de Precisión, Instituto de Medicina Oncologica y Molecular de Asturias, 33193 Oviedo, Spain

**Keywords:** Inner ear, outer hair cell, stereocilia stiffness, radixin, physiology, bundle mechanics, electrical potentials, mechanical stability, amplification, hearing loss

## Abstract

The stereocilia of the sensory cells in the inner ear contain high levels of the actin-binding protein radixin, encoded by the *RDX* gene. Radixin which is associated with mechanotransduction process such as PIP_2_ is known to be important for hearing but its functional role remains obscure. To determine how radixin influences hearing sensitivity, we used a custom rapid imaging technique to directly visualize stereocilia motion while measuring the amplitude of the electrical potentials produced by sensory cells during acoustic stimulation. Experiments were performed in guinea pigs, where upon blocking radixin, a large decrease in sound-evoked electrical potentials occurred. Despite this decrease other important functional measures, such as electrically induced sensory cell motility and the sound-evoked deflections of stereocilia, showed a minor amplitude increase. This unique set of functional properties alterations demonstrate that radixin is necessary to ensure that the inner ear converts sound into electrical signals at acoustic rates. Radixin is therefore a necessary and important component of the cochlear amplifier, the energy-consuming process that boosts hearing sensitivity by up to 60 dB.

## Introduction

The sensory cells of the inner ear are equipped with stereocilia, which harbor a molecular machinery that permits sound to be converted into electrical potentials. The protein radixin appears to be an important component of this machinery, since radixin-deficient mice are deaf^1^ from an early age and mutations in the human *RDX* gene is a cause of non-syndromic neurosensory hearing loss (DFNB24; MIM #611022, ref. ^2, 3^). Because of the effect of mutations, it is clear that radixin is necessary for normal hearing, but the physiological role of the protein remains obscure.

Radixin is enriched within stereocilia^4^ and bioinformatic analyses suggest that it is a hub in a network of interacting molecules^5^ associated with the mechanotransduction process, such as phosphatidylinositol-4,5-bisphosphate (PIP2, ref. ^6^). While the functional relevance of these interactions has not been clarified, it is evident that phosphorylated radixin links the actin cytoskeleton with various transmembrane adhesion proteins, such as CD44^7, 8^.

Given radixin’s central role in the network of proteins within stereocilia, we hypothesized that radixin may contribute to the regulation of cochlear amplification. The cochlear amplifier uses force generated within the soma of outer hair cells^9^ or within their stereocilia^10, 11^ to establish normal hearing sensitivity and frequency selectivity. The underlying mechanisms are however controversial and evidence for active force generation by mammalian stereocilia remain contentious^12^, in part because the effects of stereocilia force production are difficult to experimentally separate from the effects of the somatic motor.

To determine the influence of radixin on cochlear amplification and sensory cell function, we used a custom rapid confocal imaging technique to examine stereocilia motion while recording the electrical potentials produced by the sensory cells during acoustic stimulation. These measurements revealed an unusual pattern of functional changes when radixin was disabled. The sound-evoked electrical potentials were substantially reduced despite other important functional measures, such as stereocilia deflections and electrically induced motility, being intact. This shows that radixin allows mechanically sensitive ion channels to work at acoustic rates, suggesting radixin is a component of the cochlear amplifier acting at the level of stereocilia.

We also provide a clinical characterization of patients with RDX variants. Their hearing was normal early in life, presumably because ezrin partially substitutes for radixin, but hearing was lost during the first months of life. This causes a delay in diagnosis but also means that a brief therapeutic window exists in the event that specific therapies aimed at DFNB24 become available.

## Results

### Clinical findings in patients with mutations in the RDX gene

The first patient was a 2-year old female of Moroccan origin born to term after a normal pregnancy and delivery. Maternal serology was positive for rubella and negative for hepatitis B, human immunodefiency virus, *Toxoplasma gondii*, and syphilis, ruling out these agents as contributors to congenital hearing loss. There was no risk for chromosomal abnormalities or metabolopathies, as revealed by standard screening. The only risk factor was consanguinity, as her parents were cousins. Importantly, hearing screening before the third day of life revealed that otoacoustic emissions, faint sounds produced by the inner ear in response to low-level acoustic clicks, were present. Since the sensory outer hair cells must be intact for otoacoustic emissions to be generated, this ruled out clinically significant peripheral hearing loss (see *e.g.* ref. ^13^).

However, at the age of 16 months the patient was referred to the ENT department because of suspected hearing loss. At this time, otoacoustic emissions could not be detected, suggesting that peripheral hearing loss had developed. Auditory evoked potentials were absent and steady state evoked potential testing revealed a bilateral threshold of 90 dB hearing level at 0.5 and 1 kHz (a pedigree and the patient’s evoked potential audiogram are shown in Figure 1A). These findings are diagnostic of profound hearing loss.

**Figure 1.**
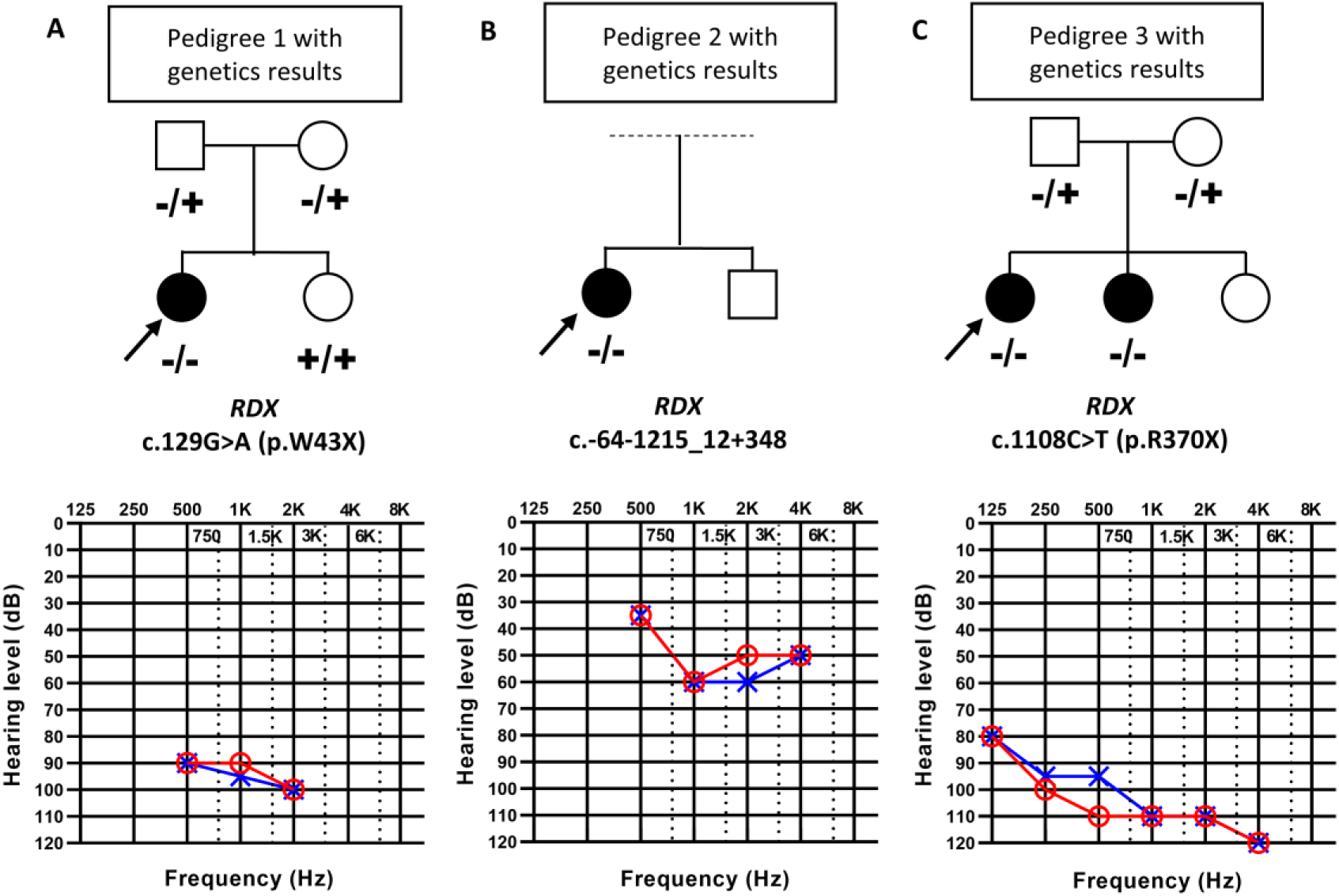
Hearing impairment in patients with *RDX* mutations. Pedigrees of the families with non-syndromic sensorineural hearing loss (a: patient I, b: patient II, c: patient III and IV). The probands are shown with arrows. Open symbols: unaffected; filled symbols: affected. Audiograms and steady state evoked potentials showed different degrees of bilateral sensorineural hearing loss of affected individuals (red, right ear; blue, left ear). (**A**) Steady state evoked potentials revealed profound bilateral hearing loss in patient I at 16 months of age. (**B**) Steady state evoked potentials showed moderate bilateral sensorineural hearing loss at 8 months of age. (**C**). Audiogram of patient III showed profound bilateral sensorineural hearing loss at 8 years of age. The variant found in this patient was included in a list of mutations in hearing-loss genes (36), but no further information about the patient was provided. Hearing thresholds of all four patients show a sloping configuration ranging from mild (patient II) to severe (patients I, III and IV) sensorineural hearing loss at low frequencies and profound impairment at high frequencies.

Genotyping with the OTOgenics panel^14^ revealed a homozygous mutation in the *RDX* gene (NM_002906.3: c.129G>A, p.W43X), which was confirmed with Sanger sequencing. The mutation truncated the protein in exon 3 (of 14), leaving only a part of the membrane-binding domain but stripping all of the actin-binding C-terminus, a change that completely disables radixin since most of its length is lost.

The second patient was female and adopted at 6 months of age. Early hearing screening was performed with brainstem auditory evoked potentials and the patient passed. However, she was referred to the ENT department at 8 months of age with a suspicion of hearing loss. Testing with steady-state evoked potentials showed moderate hearing loss (Figure 1B). Genotyping indicated a homozygous deletion of all of *RDX’s* second exon, where the initiation codon is located (NM_002906.3: c.-64-1215_12+348). Notably, an in-frame start codon present in exon 3 may mean that a protein 11 amino acids shorter is present in this patient. This shortened protein should be capable of attachment to the actin cytoskeleton, but the mutation will interfere with membrane binding.

Our third case was diagnosed with hearing loss in infancy and underwent pure tone audiometry at the age of 8 years, revealing a bilaterally symmetrical profound sensorineural hearing loss (Figure 1C). Exome sequencing disclosed a homozygous nonsense variant in exon 11 of *RDX* (NM_002906.3: c.1108C>T, p.R370X). This removes the highly conserved actin-binding motif (exons 13 and 14, ref. ^2^), preventing radixin from interacting with actin filaments. Otoacoustic emissions were absent, and a younger sister was similarly affected with symmetrical profound sensorineural hearing loss without otoacoustic emissions. Both siblings had the same homozygous *RDX* variant. Neonatal hearing screening was not performed in either case.

Overall, these clinical data show that patients with *RDX* mutations can have normal hearing on the first days of life, but hearing sensitivity deteriorates thereafter. It is not clear why this hearing loss develops, so we performed additional experiments to determine the functional role of radixin.

### Radixin expression in the hearing organ

To study radixin’s influence on hearing and the role of the protein for stereocilia function (Figure 2A), we used temporal bone preparations isolated from guinea pigs (Figure 2B), a species with low-frequency hearing similar to humans. In these isolated preparations, which retain the passive mechanics of the hearing organ^15^, direct visualization of sound-evoked stereocilia motion is possible in a nearly native environment (Figure 2C and D, ref. ^16^), which makes the preparation useful for investigating functional changes in stereocilia. However, the presence and distribution of radixin has not previously been examined in guinea pig hair cells, so we began by staining the mature organ of Corti with phosphospecific antibodies targeting radixin’s threonine 564 residue.

**Figure 2.**
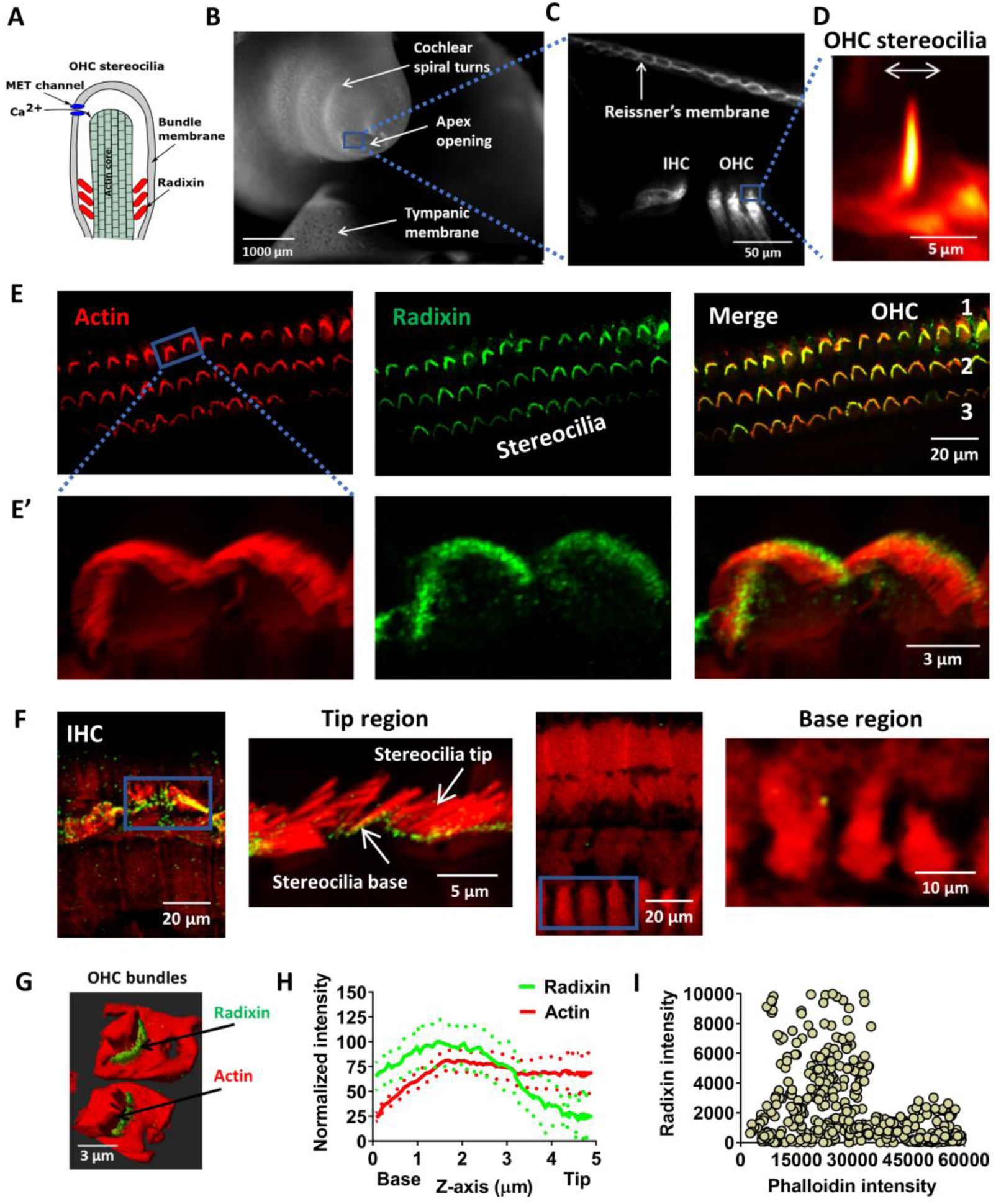
Radixin expression and localization in guinea pig cochlear hair cells. . (**A**) Schematic diagram showing the putative function of radixin in stereocilia. (**B**) A low magnification image of the temporal bone preparation. Note the apical opening used for imaging. (**C**) Release of the dye di-3-ANEPPDHQ into the endolymphatic space stained Reissner’s membrane as well as the hair bundles. (**D**) Outer hair cell (OHC) stereocilia imaged during sound stimulation at 220 Hz, 80 dB SPL. (**E**) Representative confocal images of sections of the organ of Corti labelled with a radixin-specific monoclonal antibody (green) as well as phalloidin (red, staining actin), and overlay. The bundles of the sensory hair cells are intensely labeled by the radixin antibody. OHC 1, 2, 3 indicate the three rows of outer hair cells. Images were taken from the surface preparations of the apical turn. (**E**’) Inset showing a higher magnification view. (**F**) Three-dimensional reconstruction of the organ of Corti. A close-up on the inner hair cell area shows absence of radixin label in the cell bodies of the inner hair cells. Likewise, no radixin label was detected in the neuronal or synaptic region of the inner hair cells (right side). (**G**) A 3D reconstruction of outer hair cell stereocilia showing predominance of radixin labeling near the stereocilia base and consistent actin labeling in the hair cell body and stereocilia bundles. (**H**) Normalized average signal intensity profiles for radixin and actin expression (average of 11 bundles from 3 different animals) which shows decline in radixin labeling toward the tip of stereocilia and consistent actin labeling by phalloidin throughout the stereocilia. (**I**) Scatter plot showing lack of relation between radixin and phalloidin (staining actin) pixel intensities. A.u., arbitrary units.

Immunofluorescence was observed in the stereocilia of the sensory outer and inner hair cells, with the strongest labeling in the three rows of outer hair cells. Double-labeling with fluorescently tagged phalloidin, which binds actin filaments (Figure 2E, left column), was used to locate stereocilia. These were intensely labeled by radixin antibodies (Figure 2E, center and right columns), whereas no consistent radixin label was present in either the cell bodies of the sensory cells, in their adjacent supporting cells, or in the synaptic regions of the inner hair cells (Figure 2F).

Three-dimensional reconstructions of stereocilia (Figure 2G) showed that radixin labeling was most intense in the mid-basal part of stereocilia and tapered off toward their tip. To quantify this more precisely, we measured the fluorescence intensity of each probe as a function of distance from the base of the hair bundle. Plots of the normalized fluorescence profiles (Figure 2H) confirmed the stronger labeling near the base of stereocilia, unlike the actin probe (phalloidin), which had similar labeling intensity through the length of the stereocilia.

Since the actin probe had stronger emission, we were concerned that its fluorescence might bleed through into the radixin channel. If this were the case, a linear relationship between their fluorescence intensities is expected. However, no such relationship was found (Figure 2I). This pattern of labeling is consistent with the one found in chick^17^ and rat^18^ inner ears, so we conclude that guinea pigs are an adequate model for investigating the functional role of radixin in the mature hearing organ. Next, we performed physiological measurements by combining rapid confocal imaging of sound-evoked stereocilia motion with electrophysiology, measurements of electrically evoked motion, fluorescence recovery after photobleaching, and *in vivo* measurements of hearing sensitivity in animals treated with radixin inhibitors.

### Radixin influences stereocilia deflections

Having established that radixin is present in guinea pig hair cells, but not detectable in supporting cells or in afferent neurons, we proceeded by examining the sound-evoked responses of stereocilia. To label stereocilia, a double-barreled glass microelectrode with 3-μm tip diameter was positioned close to the sensory cells. One electrode barrel was used for introducing the fluorescent dye di-3-ANEPPDHQ, which stained stereocilia (Figure 2C) and allowed their sound-evoked motion to be studied using time-resolved confocal imaging ^19, 20^. The other electrode barrel was used for delivering the radixin blocker DX-52-1, which disrupts radixin’s ability to link the actin cytoskeleton with the cell membrane (ref. ^21, 22^; Figure 3A). The loss of these interactions creates an effect similar to the truncating mutations described in our patients.

**Figure 3.**
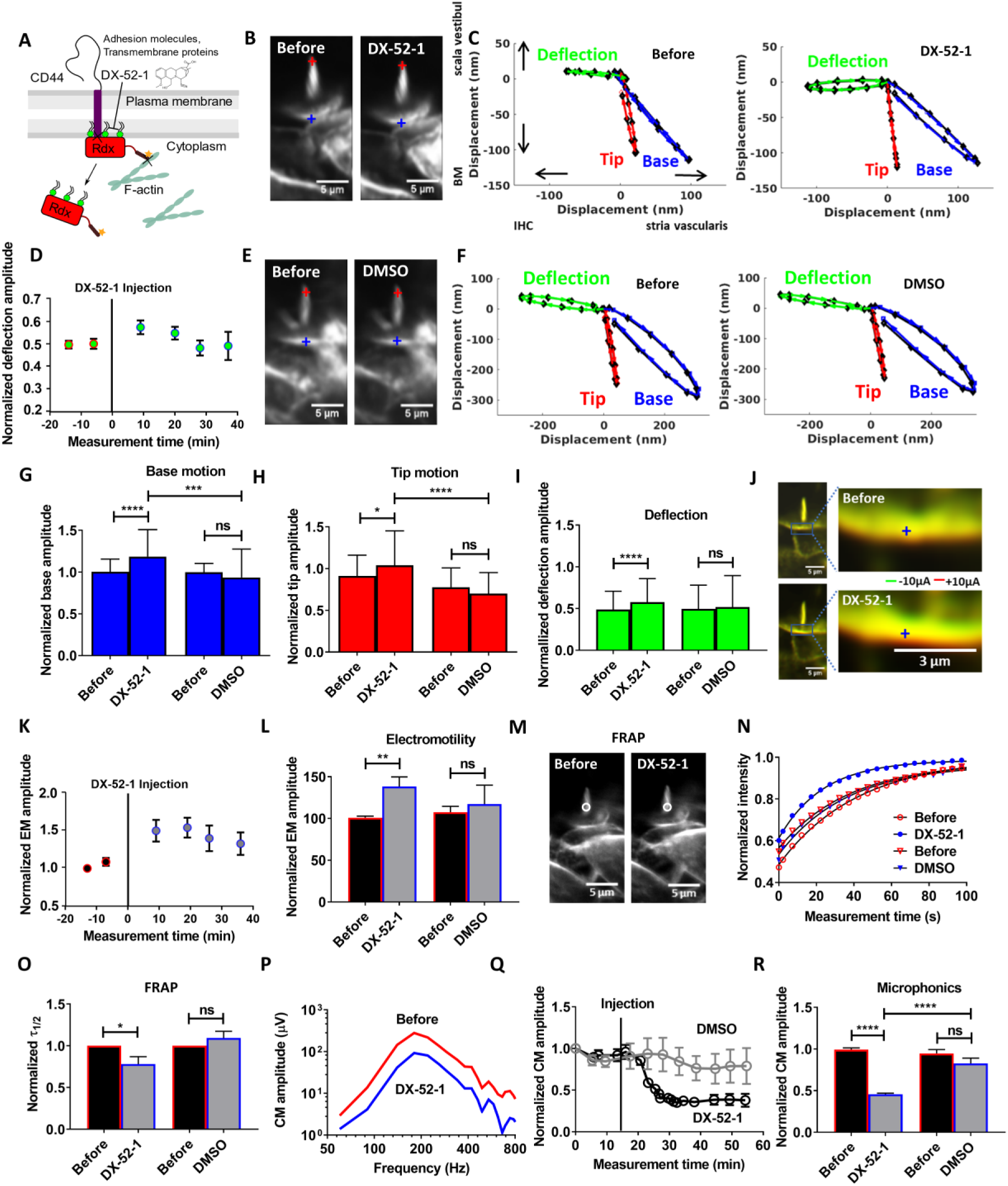
DX-52-1 induced effects in the OHC stereocilia functions. (**A**) Schematic showing how DX-52-1 disrupts radixin’s ability to interact with both actin, cell adhesion molecules and transmembrane proteins. (**B**) Time-resolved confocal images acquired during sound stimulation showing the morphology of the OHC bundle is intact before and after drug injection, except for a small change in the brightness of the fluorescent dye. (**C**) Sound-evoked motion of the bundle tip (red) and base (blue) before (left) and after (right) DX-52-1 injection in an example preparation. The stimulus was a pure tone at 220 Hz and 80 dB sound pressure level. By subtracting trajectories from the tips and bases of stereocilia, a measure of the deflection of the bundle (green) is obtained. (**D**) Time course of the averaged deflection amplitude of outer hair cell bundles (blue circle). The vertical line at time zero indicates the time of injection. Data were normalized to the average trajectory amplitude recorded before injection. Averaged data from 70 individual preparations ± standard error of the mean. (**E**) Confocal image obtained after DMSO injection, showing lack of effect on stereocilia morphology. (**F**) No change in the motion of the bundle tip (red) and base (blue) before and after DSMO injection observed along with absence of change in deflection (green). (**G** - **I**) Averaged bundle motion change at the base of outer hair cell stereocilia (blue bar), at their tip (red bar) and the deflection of the bundle (green bar). Data were normalized to the base trajectory amplitude recorded before the injection. AverageddData from 70 individual preparations ± s.d. (**J**) An OHC stereocilia bundle showing change in electrically evoked motility. Images before and after DX-52-1 were superimposed. (**K**) Time course of the averaged electromotility amplitude showing increase after DX-52-1 injection. The vertical line at time zero indicates the time of injection. Data were normalized to the average electromotility amplitude recorded before injection. (**L**) On average the electromotility amplitude increased significantly after the DX-52-1 injection (n=70) with no significant change after DMSO injection (n=15). The acoustic stimulus is a 220Hz tone at 80 dB with current stimulus of 10 μA. (**M**) FRAP experiment showing no change in the stereocilia bundle morphology before and after DX-52-1 injection, except for slight change in the dye intensity. (**N**) Normalized traces of the fluorescence intensity showing change in the membrane dynamics during the fluorescence recovery in the bundle region of interest measuring the diffusion time of the dye before and after injection averaged across 15 preparations. (**O**) Fitting the experimental data to single phase exponential fit model showed a significantly faster recovery of bundle fluorescence with reduced *τ*_*1/2*_ after DX-52-1 injection (n=24) with no change in the diffusion time after DMSO injection (n=14). (**P**) Tuning curves for the cochlear microphonic potential (CM) before and after 1.0 mM DX-52-1 injection in an example preparation. The amplitude of the CM decreased by 138 μV at its peak. (**Q**) Averaged time courses of the normalized mean peak amplitude of the cochlear microphonic potential which decreased substantially 10-15 minutes after DX-52-1 injection (n=40) but not significantly after DMSO (n=11). The vertical line indicates the time of injection of DX-52-1 and DMSO. (**R**) Comparison of the CM amplitude which reduced significantly before and after DX-52-1 injection but not after DMSO injection for experiments in panel **Q**. A significant difference in the microphonic amplitude was observed between DX-52-1 and DMSO. All data sets were normalized to the data recorded before injection. Data are the means ± s.e.m or s.d. (**G**-**I**). ****P<0.0001; ***P<0.001; **P<0.01; *P<0.05; n.s., not significant; two-tailed paired t test, two-tailed unpaired t test with Welch’s correction.

After injecting a 1-mM solution of DX-52-1 dissolved in artificial endolymph, no morphological changes were observed in stereocilia (except for minor alterations in the brightness of the dye, Figure 3B; note that the effective inhibitor concentration is reduced because the injected solution is dissolved in the endolymph present in scala media), but the injection changed the response to acoustic stimulation. Before DX-52-1 (Figure 3C, left graph), the base of the hair bundle (blue trajectory) had a different direction of motion than the bundle tip (red trajectory). As a result of this difference, motion directed at scala tympani (downwards in Figure 3C) caused deflection of stereocilia toward the center of the cochlear spiral (green trajectory). Ten to fifteen minutes after DX-52-1 (Figure 3C, right graph), sound-evoked displacement showed a minor but significant increase both at the base and at the tip of the hair bundle. As a consequence, the sound-evoked deflection of the hair bundle became larger (green trajectory in the right graph in Figure 3C). In preparations treated with vehicle alone (endolymph with 1.8% DMSO), neither morphology nor motion trajectories changed (Figure 3E and F).

The change induced by DX-52-1 was apparent 10 minutes after its application and deflections remained elevated for at least 10 minutes thereafter (Figure 3D, n=70). This period of elevated sound-induced motion was followed by gradual recovery. Figure 3G-I shows the hair bundle motion change across 70 preparations. At both the tip and the base of the hair bundle, the motion amplitude increased (from 98 ± 15 nm to 116 ± 48 nm at the base; p<0.0001, two-tailed paired t test, Figure 3G; and from 90 ± 24 nm to 102 ± 40 nm at the tip; p=0.04, Figure 3H). Base motions were larger than the tip motion, as previously described^8^. The change in the deflection amplitude was also significant (from 48 ± 21 nm to 56 ± 28 nm; p<0.0001, two-tailed paired t test; Figure 3I). A significant difference was also found when preparations injected with DX-52-1 were compared to those injected with vehicle alone (Figure 3G and h; n= 27; two-tailed unpaired t test with Welch’s correction).

In summary, the data shown in Figure 3B-I demonstrate that the radixin blocker DX-52-1 affected the sound-evoked motion of stereocilia, causing mildly increased deflection amplitudes. This finding clearly cannot explain the hearing loss seen in patients, but it is consistent with an effect of radixin on the stiffness of stereocilia.

### Radixin affects electrically evoked motility

Outer hair cells contain a transmembrane protein, prestin, which confers upon the cell the ability to rapidly change length in response to alterations in membrane potential^9^. This electromotility is critical for hearing, and to further probe radixin’s influence on hair cell function, we measured electrically evoked motility using the rapid imaging technique described above. The double-barreled microelectrode allowed us to apply 10-μA square wave currents at the frequency of 5 Hz. These currents changed the electrical potential in scala media, resulting in increased currents through the MET channels and increased force production by outer hair cells^23^.

To show the change in electromotility evoked by DX-52-1, Figure 3J shows an outer hair cell imaged *in situ* during electrical stimulation. The green channel was acquired during the negative part of the square wave and the red channel during its positive phase. Before DX-52-1 application, most pixels overlapped, signifying low motility amplitude (Figure 3J, top right graph). After DX-52-1 was introduced, the green and the red channels separated, implying an increased amplitude of electromotility (Figure 3J, bottom right graph). These changes were quantified through optical flow analysis. The time course (Figure 3K) shows that the increase was evident 10 minutes after injection of the blocker and that the amplitude remained elevated during 20 – 25 minutes. A tendency to recovery was seen thereafter. Overall, the change induced by DX-52-1 was statistically significant (from 101 ± 2 nm to 139 ± 12 nm; p=0.001, two-tailed paired t test; n=70), but this was not the case in preparations injected with the vehicle alone (Figure 3L), where no change in motion occurred.

Electrically evoked motility requires current passing through stereocilia and into the cell bodies of the outer hair cells, as evidenced by decreased amplitudes of electromotility when mechanically sensitive ion channels were blocked. Since we found an increased amplitude of electrically evoked motion, these channels must still be able to pass current.

The increase in electromotility is consistent with a slightly decreased organ of Corti stiffness, in agreement with the changes in sound-evoked stereocilia motion described above. However, neither finding explains why hearing is lost in patients with *RDX* mutations.

### The site of action of DX-52-1 is the stereocilia

To verify that DX-52-1 acts at the level of the stereocilia, we exploited the fact that radixin connects the cell membrane with the underlying actin cytoskeleton. Hence, inhibition of radixin is expected to remove an obstacle to diffusion, increasing the mobility of membrane lipids. Lipid mobility can be measured using fluorescence recovery after photobleaching (FRAP)^24^. In brief, a laser beam was focused to a submicron spot to bleach a region of interest on the stereocilia (Figure 3M). Since diffusion will add new dye molecules to the bleached area, the gradual recovery of fluorescence provides a measure of lipid mobility in the membrane, as seen in the graph in Figure 3N. Here, a single-phase exponential model (black line) was fitted to the averaged fluorescence recovery curve measured before (red open circles) and 10 - 15 minutes after DX-52-1 injection (blue circles). The fit parameters revealed significantly faster fluorescence recovery during the 25-30 minutes that followed inhibition of radixin (Figure 3O; 22 ± 2 s vs. 14 ± 1 s; p=0.02, two-tailed paired t test; n= 24). Control injections in 14 preparations showed no significant change in the fluorescence recovery time (Figure 3N). The normalized diffusion time was slightly longer after vehicle injection (Figure 3O), but this change was not significant.

The changes in lipid mobility are consistent with disruption of membrane – cytoskeletal interactions when radixin is blocked.

### Radixin inhibition decreased cochlear microphonic potentials

During sound stimulation, ions permeate mechanically sensitive ion channels from the surrounding fluid, generating extracellular electrical potentials that can be measured through the electrode placed near the sensory cells. By tracking the amplitude of these microphonic potentials over a range of stimulus frequencies, tuning curves were acquired.

Upon injection of DX-52-1, a decrease in the cochlear microphonic (CM) amplitude (Figure 3P) was evident 10-15 minutes after the blocker injection, and the amplitude remained depressed during the ensuing 30 – 35 minutes (Figure 3Q, n=70). On average, the CM amplitude decreased from 124 ± 16 μV to 57 ± 9 μV, measured at the peak of each tuning curve (Figure 3R, p<0.0001, two-tailed paired t test). A significant difference in the amplitude was also evident between preparations injected with DX-52-1 and the controls (p<0.0001, two-tailed unpaired t test with Welch’s correction; n= 13 controls).

The decrease in the CM amplitude means that the ability to convert sound into rapidly alternating electrical potentials is impaired. This however is not due to a change in the stimulation of stereocilia, because stereocilia deflections were slightly increased (Fig. 3B-I). Also, the decrease in the CM is not due to a blocking effect on mechanically sensitive channels, as demonstrated by the increase in electrically evoked motion (Fig. 3J-L), which requires currents to pass through these channels into the hair cell soma.

### Compound action potentials indicate loss of hearing sensitivity in vivo

To assess the influence of radixin on hearing sensitivity *in vivo*, we applied 1 μl of a 1 mM DX-52-1 solution directly to the round window membrane of anesthetized guinea pigs while measuring the amplitude of the auditory nerve compound action potential (CAP). The CAP represents the summed response of auditory nerve fibers to acoustic stimulation, and is most effectively elicited by high-frequency acoustic stimuli with rapid rise time. Ten to 40 minutes after the application of DX-52-1, the CAP amplitude decreased significantly compared to control preparations where only the vehicle, perilymph with 1.8% DMSO, was applied (Figure 4A).

**Figure 4.**
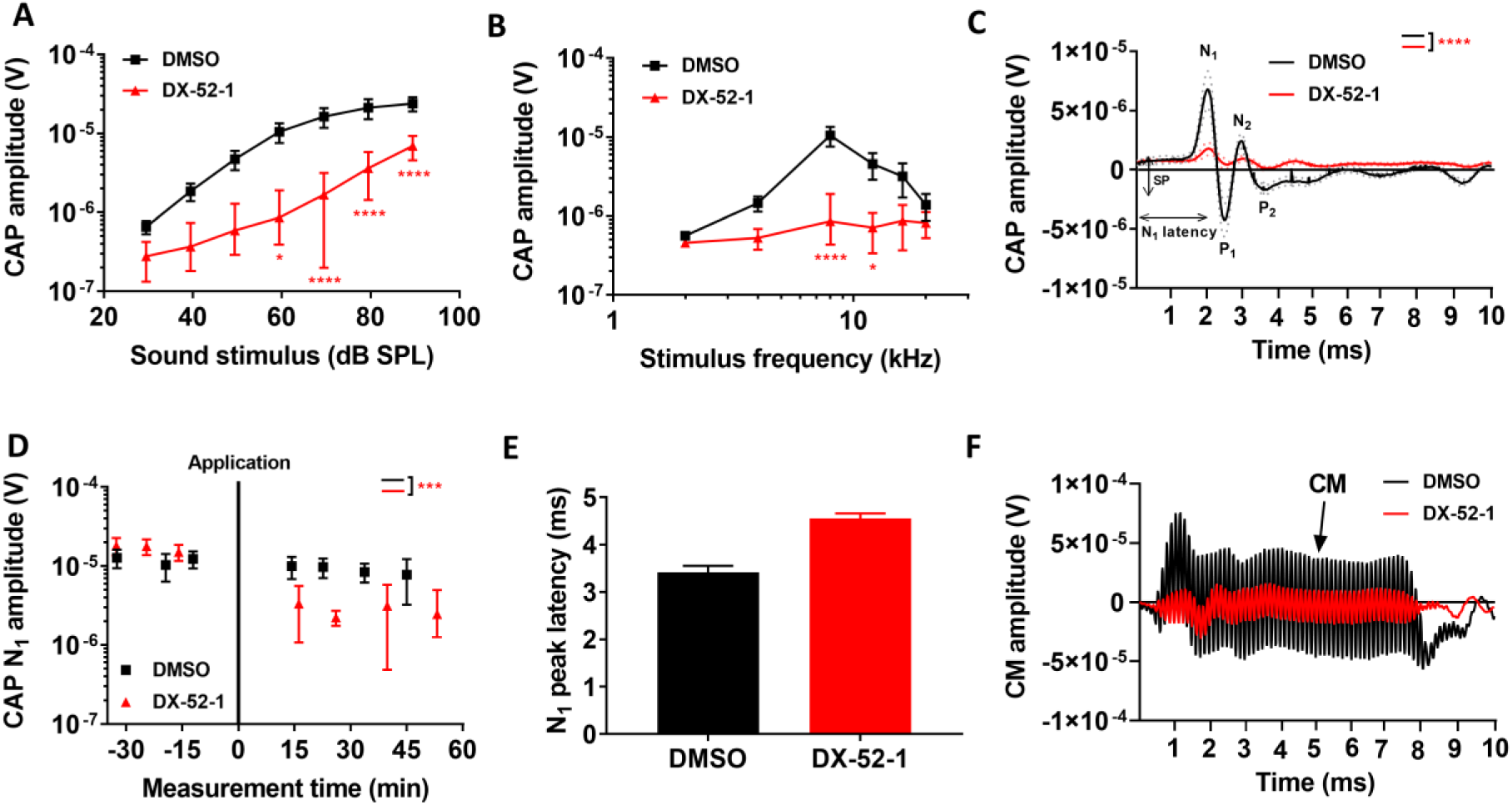
DX-52-1 results in declining hearing sensitivity, as assessed by the compound action potential of the auditory nerve. (**A**) Schematic showing the CAP recordings for control (black) and DX-52-1(red) treated guinea pigs. (**B**) Average CAP amplitude to 60 dB SPL stimuli shows a reduction for the DX-52-1 animals compared to control. (**C**) Grand averages ± s.e.m (dotted) of the CAP waveforms to 60 dB SPL 8-kHz stimuli shows reduction in N1 and N2 amplitudes. (**D**) Averaged time courses of the changes seen in N1 amplitude measured at 60 dB SPL 8-kHz stimulus following DX-52-1 application, relative to those before the application, which decreased significantly after 15-20 minutes of application. (**E**). Comparison of the CAP N1 latency which increased slightly after DX-52-1 application for animals in panel **C**. (**F**) Representative waveforms of the cochlear microphonic potential (CM), reflecting OHC activation before and after 20 minutes of application of 1 mM DX-52-1. The stimulus was a 8-KHz tone burst at 90 dB SPL. The vertical line at time zero indicates the time of application. Data information: DMSO (n=10), DX-52-1 (n=18). ****P<0.0001; ***P<0.001; **P<0.01; *P<0.05; ns, not significant; two-way ANOVA coupled to the Bonferroni post hoc test, two-tailed unpaired t test with Welch’s correction. Data are the means ± s.e.m.

Analysis of CAPs confirmed that hearing impairment was most pronounced at frequencies between 8 and 16 kHz, while smaller changes were observed at other frequencies (Figure 4B; n=18 for DX-52-1 vs 10 controls; p<0.0001; two-way ANOVA). While the overall shape of the CAP waveform remained similar after DX-52-1, there was a slight increase in the response latency (Figure 4C, E). Figure 4D demonstrates the time course for the change in CAP N1 peak amplitude, with maximum amplitude change after about 20 - 30 min. As shown in Figure 4F, DX-52-1 decreased the amplitude of the cochlear microphonic potential (in Figure 4F, the stimulus was a 90-dB SPL tone at 8 kHz).

The above data show that inhibition of radixin has effects that parallell the human data, where *RDX* mutations caused profound hearing loss.

### PAO-induced effects on stereocilia function

Radixin mediates interactions between the cytoskeleton and the cell’s membrane, but membrane attachment also requires the presence of PIP2, the synthesis of which can be blocked by kinase inhibitors such as phenylarsineoxide (PAO; Figure 5A and ref. ^6^). Although the rates of both fast and slow adaptation are affected by PAO^6^, its indirect inhibitory effect on radixin can be used to confirm some of the DX-52-1 effects described above.

**Figure 5.**
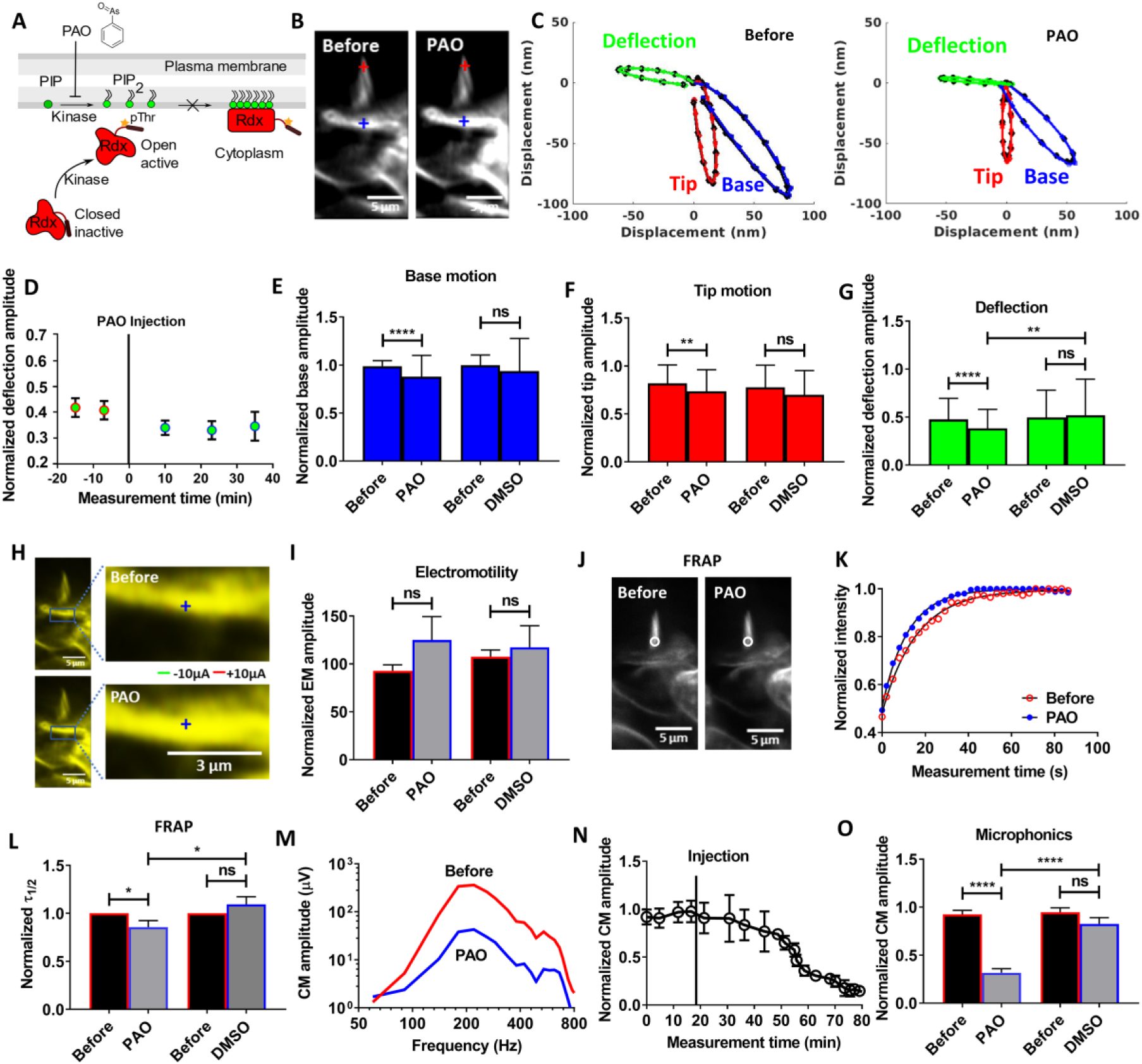
PAO induced effects in the OHC stereocilia functions. (**A**) A Schematic demonstrating the mechanism for the regulation of radixin protein via PIP2 binding. PAO inhibits the synthesis of PIP2 by blocking PI4 kinase, thus decreasing the levels of PIP2 and preventing activation of radixin. (**B**) Time-resolved confocal image of an OHC stereocilia bundle showing the morphology is intact before and after the injection, except for a small change in the brightness of the fluorescent dye. (**C**) Representative data showing change in sound-evoked motion of the bundle tip (red) and base (blue) before and after PAO injection. (**D**) Time course of the averaged deflection amplitude of outer hair cell stereocilia bundle (blue circle) showing decrease after PAO injection. The vertical line at time zero indicates the time of injection of PAO. Data were normalized to the average trajectory amplitude recorded before injection. Averaged data from 35 individual preparations ± s.e.m. (**E**-**G**) Averaged change of the bundle motion at the base of outer hair cell stereocilia (blue bar), at their tip (red bar) and deflection (green bar) shown. Significant decrease in tip and base motion resulting in a change in the bundle deflection. Data were normalized to the base trajectory amplitude recorded before the injection. Mean data from 35 individual preparations ± s.d. (**H**) An OHC bundle showing electrically evoked cell motility change color-coded displacement data superimposed before and after PAO injection. (**I**) The average electromotility amplitude increased non-significantly 30 nm after the PAO injection. Data from 28 individual preparations. The acoustic stimulus is a 220 Hz tone at 80 dB with current stimulus of 10 μA. (**J**) No change in the stereocilia bundle morphology seen before and after PAO injection for FRAP experiment. (**K**) Normalized traces of the fluorescence intensity during fluorescence recovery in the bundle region of interest measuring the diffusion time of the dye before and after injection averaged across 22 preparations. (**L**) Fitting the experimental data to single phase exponential fit model showed a faster recovery of the bundle fluorescence with reduced *τ*_*1/2*_ after PAO injection on average for experiments in panel **N**. (**M**) Tuning curves for the cochlear microphonic potential before and after 1.0 mM PAO injection in an example preparation. The amplitude of the cochlear microphonic potential decreased by 320 μV at its peak. (**N**) Normalized mean peak amplitude of the averaged time courses of the cochlear microphonic potential showing substantial irreversible decrease 10-15 minutes after PAO injection (n=33). The vertical line indicates the time of injection of PAO. (**O**) Comparison of cochlear microphonic potential amplitude before and after PAO injection which reduced significantly for experiments in panel n. Data were normalized to the data recorded before the injection. Data are the means ± s.e.m or s.d. (**E**-**G**). ****P<0.0001; **P<0.01; *P<0.05; n.s., not significant; two-tailed paired t test.

**Figure 6.**
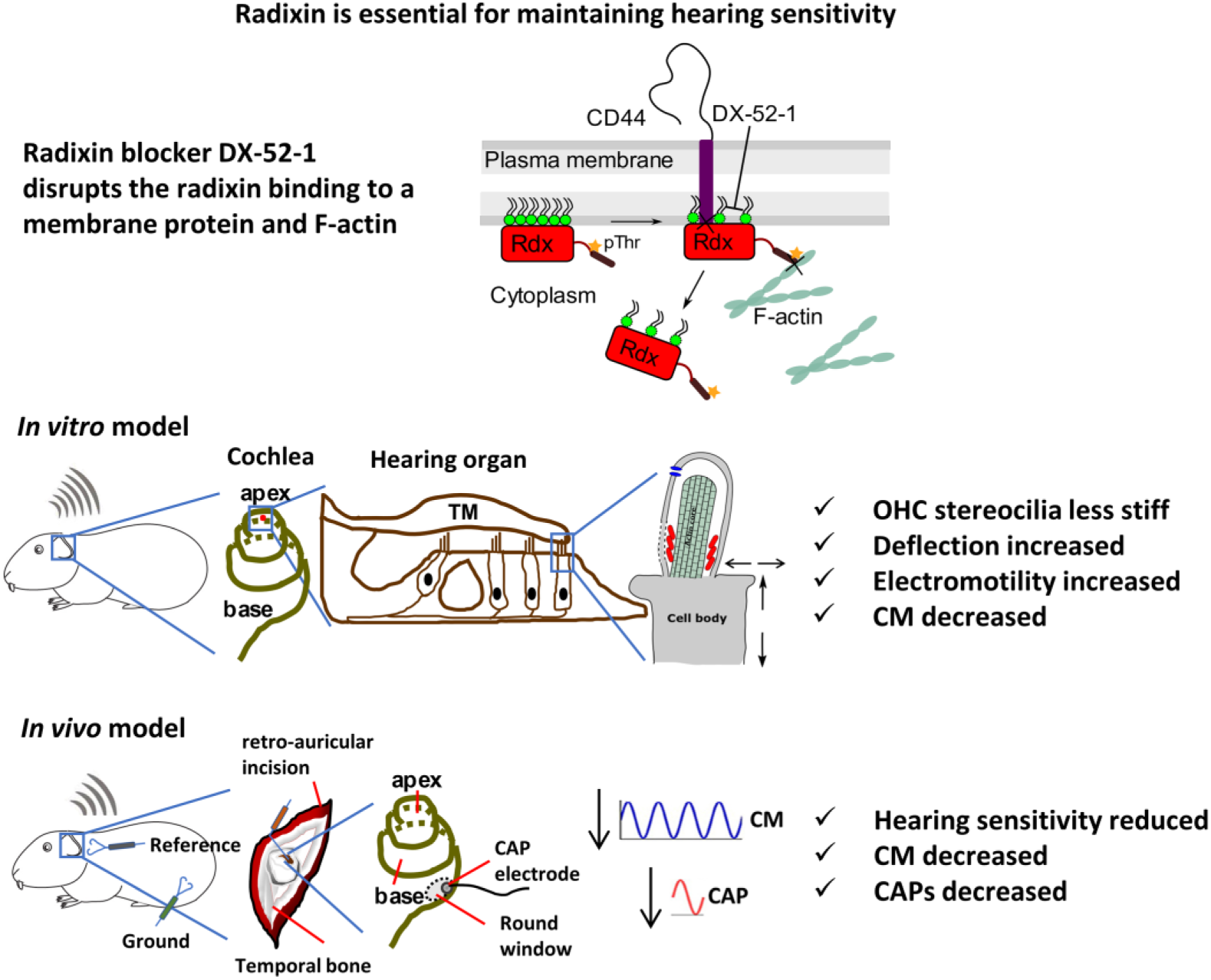
Radixin is required for maintaining the mechanical stability of stereocilia and hearing sensitivity. Schematic diagram of outer hair cell stereocilia with radixin binding area. The top scheme represents the molecular interactions between radixin and F-actin cytoskeleton and the transmembrane protein CD44. In the hearing organ of animals where the radixin blocker DX-52-1 was not applied, the animals had normal hearing and stereocilia functions. Application of the blocker results in a disruption of the link between radixin and F-actin. The animal had reduced hearing sensitivity and large effects on the OHC stereocilia functions were evident.

Injection of a 1-mM PAO solution into the endolymphatic space produced minor changes in brightness of outer hair cell stereocilia, but no other morphological changes were evident (Figure 5B). As seen in the example data in Figure 5C, the sound-evoked displacement at both the tip of the stereocilia (red trajectory) and at their base (blue trajectory) decreased following PAO. This decrease led to a reduced deflection amplitude (green trajectory; right panel in Figure 5C), even though the shapes of the motion trajectories remained similar. The change in deflection amplitude was apparent 10 – 15 minutes after PAO injection and the amplitude continued to be reduced over the ensuing 40 minutes (Figure 5D, n=35).

Aggregated data across 35 preparations are shown in Figure 5E - F. The decrease in motion amplitude at the base of stereocilia was significant (from 97 ± 6 nm to 86 ± 22 nm; p<0.0001, two-tailed paired t test; Figure 5E) as were the change in displacement at their tips (from 80 ±19 nm to 72 ±22 nm; p<0.004, 2-tailed paired t test; Figure 5f). The deflection amplitude decreased from 46 ±21 nm to 37 ± 20 nm (p<0.0001, two-tailed paired t test; Figure 5G). A significant difference was also found when preparations injected with PAO were compared to those injected with vehicle alone (Figure 5G; p=0.004, two-tailed unpaired t test with Welch’s correction).

Considering that DX-52-1 caused an increased motion amplitude in response to electrical stimulation, we proceeded by examining the influence of PAO on electromotility. Color-coded data from an example preparation are shown in Figure 5H. In this case, images acquired before and after PAO largely overlapped as demonstrated by the yellow color in Figure 5H, indicating that PAO did not change electrically evoked organ of Corti motion. Across 28 preparations, there was an increase from 93 ± 6 nm to 125 ±24 nm in the mean amplitude of electrically evoked motion, but this change was not significant (p=0.20, two-tailed paired t test; Figure 5I). Also, there was no significant difference between preparations injected with the vehicle alone and those injected with PAO (Figure 5I).

Next, we used FRAP to look for changes in the membrane lipid diffusion kinetics after PAO injection. Diffusion of di-3-ANEPPDHQ molecules within a defined ROI (Figure 5J) on the stereocilia was measured. In the data shown in Figure 5K, a single-phase exponential model (black line) was fitted to the averaged fluorescence recovery curve before (red open circle) and 10 minutes after PAO injection (blue filled circles). The fit parameters revealed significantly faster (from 21 ± 3 s to 16 ± 2 s; p=0.04, two-tailed paired t test; n=22; Figure 5L) diffusion during the ensuing 25 -30 minutes. A significant difference in the fluorescence recovery time was also seen between preparations injected with PAO and the controls (Figure 5L; p=0.03, two-tailed unpaired t test with Welch’s correction).

PAO injection also led to a decrease in the CM amplitude (Figure 5M). The drop in the CM amplitude was evident within 10-15 minutes after the injection, and there was no recovery during the ensuing 30 – 40 minutes (Figure 5N; n=33). On average, the CM amplitude decreased from 145 ± 20 μV to 58 ± 10 μV, measured at the peak of each tuning curve (Figure 5O, p<0.0001, two-tailed paired t test). A significant difference in the amplitude was seen between preparations injected with PAO and the controls (Figure 5O; p<0.0001, two-tailed unpaired t test with Welch’s correction).

The change in lipid mobility evoked by PAO and the decrease in the CM amplitude are consistent with the DX-52-1 findings; however PAO is unspecific (25) and will affect many proteins found in stereocilia, which likely explains why the effects on sound-evoked motion and on electromotility differ from those of DX-52-1.

## Discussion

This study shows that radixin allows stereocilia to generate electrical potentials at acoustic rates, making radixin necessary for cochlear amplification. The effects of radixin inhibition are not due to a change in the stimulation of the sensory cells, since stereocilia deflections showed a minor increase upon blocking radixin (Fig. 3). Similarly, the decrease in the electrical potentials produced by the sensory cells is not due to inhibition of electromotility. The increase in both sound-evoked deflections and in electromotility are however consistent with a decrease in stereocilia stiffness.

Previous studies showed that hair cell stereocilia contain high levels of radixin ^1,4^,5. Some studies also demonstrated radixin labeling at the junctions between the supporting cells and the hair cells^26^, but this was not evident in our experiments and no consistent labeling was found in either neurons or in the cell bodies of the sensory cells. These results suggest that radixin inhibition affects stereocilia function. This view is supported by findings from radixin knockout mice, which show degeneration of stereocilia after the onset of hearing, but an otherwise normal organ of Corti structure^1^. It appears that upregulation of ezrin, a protein closely related to radixin, ensures normal early development of stereocilia but this compensation mechanism subsequently fails. Hence, it is clear that radixin is critical during the final phases of stereocilia development, but it continues to be expressed at high levels through the life of the animal^1^ suggesting an important physiological role that has remained obscure.

Membrane-associated proteins such as radixin are often regulated by membrane lipids. Radixin is activated only after positive regulation, which requires sequential binding of PIP2 and phosphorylation of threonine 564^27^. In hair cells, radixin is concentrated towards the stereocilia base, where they insert into the cuticular plate. This taper region is a site of mechanical stress during sound-evoked deflection^28^. Based on the findings of the present study we propose that radixin, in addition to its role for channel function, contributes to the regulation of stereocilia stiffness by linking the cytoskeleton more tightly to the membrane inside this high-stress region. Findings evident after the inhibition of radixin and consistent with this hypothesis include the increased lipid mobility (Figure 3N, O), larger electrically evoked motility (Figure 3K, L), and larger sound-evoked stereocilia deflections (Figure 3C, G-I) evident after inhibition of radixin. Due to the active, nonlinear mechanisms that amplify sound-evoked motion *in vivo*^29, 30^, small changes in the mechanical properties of stereocilia can have large effects on hearing organ performance.

However, the most dramatic effect of radixin inhibition was the reduction in sound-evoked electrical potentials and in the amplitude of the CAP. This demonstrate a previously unrecognized role of radixin in maintaining the amplitude of the mechanoelectrical transduction current. Since we (Figure 2) could detect no radixin expression either in cochlear neurons or at the synaptic pole of the hair cells, the reduction of the CAP amplitude is explained by an effect on the transduction process itself. This finding is supported by the normal morphology of the organ of Corti in aged radixin knockout mice^1^, and with the absence of detectable radixin expression in cochlear neurons^2^.

It is interestering that two of our patients had apparently normal hearing at birth, as shown by them passing the newborn hearing screening program (the 2 siblings from pedigree 3 did not undergo neonatal hearing screening). The subsequent development of hearing loss could be due to a combination of reduced transduction currents and an inability to maintain stereocilia structure, including their stiffness, in the long term in the absence of membrane-cytoskeletal links. However, hearing loss was profound in three of our patients and moderate in one. At first sight, the removal of the start codon in exon 2 in this patient should lead to complete absence of radixin expression. Due to an in-frame start codon present in exon 3, it is however possible that a protein 11 amino acids shorter could be produced. We speculate that such a shorter protein could retain some functionality, explaining the less severe hearing loss in this patient and suggesting a clinically relevant genotype-phenotype correlation for pathogenic *RDX* variants. Moreover, in one of our families copy number variations contributed to the development of the hearing loss. It is important to identify such variation during genetic testing, since it is a challenge to conventional next-generation sequencing technologies.

The development of early onset hearing loss in children that passed newborn hearing screening can confound both patients and their physicians causing diagnosis and intervention to be postponed^31, 32, 33^. The resulting delays in speech and language development may contribute to impairment of social skills and cognition^34^. No previous study has examined the effect of radixin on stereocilia function. Therefore, understanding the physiopathology of genes such as *RDX* and increasing our awareness of its contribution to this burden of delayed diagnosis could improve the care of children with hearing impairment. This is important, specially for siblings of already diagnosed patients. Therefore, in these families, if a genetic diagnosis has not been obtained, close monitoring of the siblings that have passed initial newborn hearing screening is mandatory. Importantly, the fact that hearing appeared normal early in life could mean that a time window exists in the event that therapies for restoring radixin functionality become available. In any case, those potential, gene-specific, therapeutic opportunities will always be enhanced by an early and comprehensive genetic diagnosis.

## Materials and Methods

### Ethics statement

The clinical data collection was approved by the Institutional Review board at the University of Miami (USA) and by the by the Comité de Ética de Investigación del Principado de Asturias (research project #75/14), Spain. A signed informed-consent form was obtained from each participant or, in the case of a minor, from the parents.The Regional Ethics Board in Linköping approved all animal experiments (DNR 16-14) and animal care was under the supervision of the Unit for Laboratory Animal Science at Linköping University.

### Clinical study

Patients I and II (Figure 1) were evaluated according to standard newborn hearing screening protocols using otoacustic emissions and/or auditory evoked potentials. Later, patients I and II were studied again because of a suspicion of hearing loss. Objective measures of hearing was used to establish their audiograms. In patient III, sensorineural hearing loss was diagnosed via standard audiometry in a soundproof room according to current clinical standards as recommended by the International Standards Organization (ISO8253-1). Routine pure-tone audiometry was performed with age-appropriate methods to determine hearing thresholds at frequencies 0.25, 0.5, 1, 2, 4, 6 and 8 kHz. Severity of hearing loss was determined from pure tone averages calculated at 0.5, 1.0, 2.0 and 4 kHz. Transient evoked otoacoustic emissions were tested. DNA was isolated from whole blood of the probands and panel (patients I and II) or exome (patient III) sequencing was performed as previously described^14,36^. Validation and segregation testing of the variants was performed.

### Animal and experimental model details

Young mature Dunkin-Hartley guinea pigs of both sexes (250 - 450 g) were used for all experiments. Prior to decapitation all animals were tested for the Preyer reflex and then anesthetized with 18 – 24 mg of sodium pentobarbital intraperitoneally, according to their body weight. The left temporal bone was excised from the guinea pigs and attached to a custom-built holder. The holder allowed immersion of the cochlea and the middle ear in oxygenated (95 % O_2_, 5 % CO_2_) cell culture medium (Minimum Essential Medium with Earle’s balanced salts, SH30244.FS Nordic Biolabs). The bone of the bulla was removed gently with bone cutters which exposed the middle ear and the basal turn of the cochlea, including the round window niche. Thereafter a small triangular or trapezoidal opening was made at the apex using a #11 scalpel blade and a hole of 0.6 mm diameter was drilled at the base of the cochlea using a straight point shaped pin. These openings allowed continuous perfusion of oxygenated tissue culture medium through an external syringe tube connected to the basal hole with a plastic microtube. Sound stimulation occurred through a calibrated loudspeaker connected to the chamber with a plastic tube. Because of the immersion of the middle ear and the opening at the apex, the effective sound pressure level was reduced by ∼ 20 dB. The values given throughout the text are corrected for this attenuation. The whole preparation was maintained at room temperature (22 - 24°C). The apical opening allowed confocal imaging of the hearing organ and permitted insertion of a double-barrel glass microelectrode filled with artificial endolymph-like solution (1.3 mM NaCl, 31 mM KHCO_3_, 23 μM CaCl_2_, 128.3 mM KCl, pH 7.4 and 300 mOsmol/kg adjusted with sucrose) into the scala media through the Reissner’s membrane (RM). This special electrode with septum is used for cochlear microphonic recordings (CM), electrical stimulation, endocochlear potential recordings (EP), bundle membrane staining and blockers delivery, as specified.

### Reagents

The following stock solutions were prepared and further diluted in artificial endolymph to the desired concentration to be used in the study. Di-3-ANEPPDHQ (D36801 ThermoFisher Scientific): 4.0 mM in pure DMSO diluted 100 times for use. Quniocarmycin analog DX-52-1 (a kind gift from the US National Cancer Institute, 96251-59-1): 22.0 mM in 50% DMSO and phosphate-buffered saline diluted to 1.0 mM for use. Note that the effective concentration in the endolymph is lower than 1 mM because the agent is diluted in the scala media fluids upon injection. Previous estimates suggest a 10x dilution factor (16). Phenylarsine Oxide (P3075-1G Sigma Aldrich): 45.0 mM in pure DMSO diluted to 1.0 mM for use.

### Confocal imaging

Samples were imaged with an upright laser scanning confocal microscope (Zeiss LSM 780 Axio Imager) controlled with the ZEN 2012 software. Outer hair cell bundle displacement movements were acquired with a 40X, 0.80 numerical aperture water immersion objective lens (Zeiss Achroplan or Nikon CFI Apo lens); immunofluorescence imaging was made with a 100X oil immersion, 1.40 NA objective (Zeiss Plan-Apochromat). Images were processed in Imagej 1.50i software, Imaris 9.2, ZEN 2012 and Matlab (R2017b, the Mathworks, Natick, MA, USA) and schemes drawn in Inkscape 0.92.3.

### Electrophysiological recordings

Hair bundles were labeled with the membrane dye di-3-ANEPPDHQ which was dissolved in endolymph solution and delivered by electrophoresis. This protocol ensured minimal dye release into the scala media and produced strong labeling of stereocilia while preserving the barrier function of Reissner’s membrane (RM). Double-barrel microelectrodes with an outer diameter of 1.5 mm were pulled with a standard electrode puller and beveled at 20 degrees to a final resistance of ∼4 – 6 MΩ. The electrodes were mounted in a manual micromanipulator at an angle of 30 degrees and positioned through the apical opening close to the RM. The RM was penetrated using a hydraulic stepping motor. Current injections were performed with a linear stimulus isolator (A395, World Precision Instruments) sending positive steady state currents of up to + 14 μA. These currents restored the normal potential around the hair bundles, leading to an increase of the currents through the MET channel, and in the force produced by the hair cells. The endocochlear potential upon penetration of RM was ∼25-30 mV. Cochlear microphonic potentials were measured with an Ix1 amplifier (Dagan Instruments) and digitized with a 24-bit A/D board (NI USB-4431, National Instruments) at 10 kHz, using custom Labview software. Tuning curves were recorded in response to a series of tone bursts at 60 dB SPL ranging from 60 to 820 Hz. The rise and fall time was 1 ms, using a Hanning window. The samples signals were Fourier-transformed and the peak amplitude plotted as a function of stimulus frequency. Before applying drugs, tuning curve measurements were repeated every five minutes for 15-20 minutes to verify that the response was stable. We therafter proceeded with other measurements, as described in the text.

### Time-resolved confocal imaging

To measure sound-evoked bundle motion, the hearing organ was faintly stained with 1 μl of dye di-3-ANEPPDHQ added in the perfusion tube. Subsequently, the sensory hair cell bundles were stained with di-3-ANEPPDHQ dissolved in the electrode solution and delivered to the hair bundles iontophoretically with a current stimulus of 3-5 μA. The preparation was stimulated acoustically near the bundles’ best frequency (180 – 220 Hz). The best frequency was selected from the highest peak of the tuning curve of the cochlear microphonic recordings. Image acquisition triggered both the acoustical and electrical stimulus. A series of 37 images was acquired; each series requiring ∼40 s for combined sound and current stimulus. Custom Labview software ensured that every pixel in the image series had a known phase both of the acoustic and electrical stimuli. To obtain images free from motion artefacts, the softwared tracked the temporal relation between the pixels and the sound stimulus. Image sequences free from motion artefacts were then reconstructed using a Fourier series approach^19, 20^, to generate a sequence of 12 images at equally spaced phases of the sine wave. Images for positive and negative current stimulation were also reconstructed at 12 equally spaced phases. These image sequences were low-pass filtered and subjected to optical flow analysis using a 2D version of the 3D algorithm described in ref. 20. To improve the signal-to-noise ratio, trajectories for all pixels in a 3×3 or 5×5 region were averaged. For combined sound and electrical stimuli, current injection switched directly from positive to negative at 5 Hz to avoid charge build-up in the scala media.

### Blocker injection

For experiments in which blockers (DX-52-1 or PAO) were injected into the endolymphatic space through the double-barrel microelectrode, one barrel of the electrode was filled with the dye di-3-ANEPPDHQ dissolved in endolymph and the other contained the blocker dissolved in endolymph. Pipettes had 1.5-3 μm tip diameter and were positioned 50-70 μm from the hair bundles The blocker was pressure-injected by a 2 pound-per-square inch pressure pulse lasting for 10 s. To verify the injection, a time series of confocal images, 60 to 100 s in length was acquired during each injection. Cochlear microphonic potentials (CM) were recorded before and at 5-minute intervals after the injection. Sets of confocal images of hair bundle displacement were recorded before (2 sets) and after injection (3-4 sets) and continued every five minutes for the next 30-40 minutes of the experiment time at a stimulus level of 80 dB SPL, 10 μA at 220 Hz best frequency. The argon laser line at 488 nm and matching beamsplitter was used. To avoid bleaching, the laser power was set to the minimum value consistent with an acceptable signal-to-noise ratio.

### Fluorescence Recovery After Photobleaching (FRAP)

FRAP was performed by outlining a region of interest on the stained stereocilia membrane. Following an acquisition of a series of 10 baseline images, a 2-μm spot on the stereocilia membrane was photobleached by focusing the laser at a maximum power into the region of interest^35^. The recovery of fluorescence was tracked by acquiring a series of 30 images at 1 or 3-s intervals over a time of 100 – 140 s. The images were 256 × 512 pixels, 12-bit pixel depth, with an integration time of 6.30 μs per pixel, and a pinole of 1.50 Airy units. Confocal images were obtained before and 10 and 20 minutes apart after the blocker injection. Statistical analysis was performed by fitting the experimental data to a one-phase decay model.

### Compound action potentials

To record compound action potentials (CAPs), animals were anaesthetized with an initial dose of an intra-muscular injection of Xylazin (0.5ml/kg) and Ketalar (0.4ml/kg). Three to four minutes after the animal was adequately anesthesized, the surgical site of the left bulla was shaved and the animal placed on a thermostatically controlled heating blanket to maintain a core body temperature of 38°C. Bupivacain (0.2ml/kg), a long-acting local anaesthetic, was administered near the surgical site before skin incision. A retroauricular incision was made in order to reach the temporal bone. Muscle and other soft tissues were dissected, and the posterio-lateral part of the auditory bulla was opened to access the round window niche. A thin Teflon-insulated Ag/AgCl silver ball recording electrode was placed in close contact with the round window membrane. The electrode wire was fixed to the temporal bone with dental cement to ensure the position of the recording electrode remained stable throughout the experiment. The animal was then placed inside a sound-proof recording booth where an Ag/AgCl electrode was inserted subcutaneously at the vertex of the skull. Cochlear compound action potentials were recorded sequentially from the left ear of the animal. Standardized input–output functions were generated by varying the intensity of stimulus (90 dB, 80 dB, 70 dB, 60 dB, 50 dB, 40 dB, 30 dB SPL in steps at 6 different frequencies 2 kHz, 4 kHz, 8 kHz, 12 kHz, 16 kHz, 20 kHz). The recorded evoked CAPs signal was then filtered (high-pass frequency 3-5 Hz, low-pass frequency 3-5 kHz) and amplified at a gain of 10 000 and stored for offline analysis. The responses to 200 repetitions of each stimulus were averaged with a sampling rate of 100 kHz. Three sets of recordings were obtained before the blocker application, and recordings were repeated at 5-min intervals for the next 40 minutes after application of the blocker. Blockers were dissolved in artificial perilymph (137 mM NaCl, 5 mM KCl, 2 mM CaCl_2_, 1 mM MgCl_2_, 5 mM D-glucose, 5 mM HEPES, pH 7.4, 300 mOsmol/kg) and introduced on the round window membrane. All recording software’s were custom written in LabVIEW.

### Surface preparation and immunofluorescence staining and imaging

Whole-mount preparations of the guinea pig organ of Corti were obtained as follows. Temporal bones were removed, the bony bulla was opened to visualize the cochlea and two small hole were made in the round window and at the apex. These openings allowed perfusion of the sensory epithelium with phosphate-buffered saline solution (PBS). The perilymphatic space was gently perfused with 4% paraformaldehyde in PBS. The sensory epithelium was exposed by carefully removing the cochlear bone, the spiral ligament, and the tectorial membrane. After washing the samples in PBS, they were permeabilized by treating with, 0.3% Triton X-100 soaked in 3% bovine serum albumin (BSA, 0332-25G, VWR), dissolved in PBS for 10 minutes at room temperature followed with one-time wash with PBS for 5 minutes. Permeabilization was followed with blocking step by incubating the samples for 2 hr in PBS containing 3% normal goat serum (NGS, 927503, BioLegend) and 3% BSA and then stained overnight at 4°C with the primary monoclonal antibody (mouse anti-radixin, ABNOH00005962-M06, Abnova) at a dilution of 1:500. Samples were then washed three times with PBS for 10 minutes each, followed by a 2 hr incubation with a mixture of the secondary antibody (goat anti-mouse Alexa Fluor^®^ 488–conjugated IgG, ab150113 Abcam) and Alexa Fluor 568 conjugated Phalloidin (A12380, ThermoFisher Scientific) at a dilution of 1:500. The antibody solutions were prepared in blocking solution. After three washes with PBS for 20 minutes each, sections were readied for surface preparations. Sections of organ of Corti starting from apex to base were carefully dissected and mounted on the glass slides prepared earlier and cover slipped with mounting media Fluorosave reagent (345789, Calbiochem). The slides were sealed and allowed to rest for ∼2 h before proceeding with imaging. Confocal images of the mounted sections were obtained in two track channel mode with MBS 488/561 excited at 488 nm for Alexa Fluor 488 fluorescence, emission range between 490-570 nm and at 561 nm for Alexa Fluor 568 fluorescence, emission range between 570-695 nm. Z-stacks were acquired at 12-bit pixel depth, 512 × 512 pixels, with an integration time of 6.30 μs per pixel, pinole of 1.0 Airy units and a spacing 1.0 or 3.0 μm per slice with 20 slices up to 10 μm in total depth.

### Fluorescence intensity quantification

Identical experimental settings and analyses were used for quantifying both radixin and phalloidin immunofluorescence. Maximum projections of confocal z-stacks were acquired and used for analysis. Organ of Corti sections were fixed, immunostained, mounted, and imaged. For background subtraction, fluorescence intensity from randomly chosen areas per preparation, lacking specific signal, were averaged and subtracted from the respective images. Hair bundles were outlined manually in ImageJ, and the average fluorescence intensity was calculated for each individual hair bundle. Individual fluorescence intensity values of a given experiment were normalized to the global average of the corresponding preparations.

### Data evaluation and statistical analysis

All experiments were repeated multiple times; the number of individual measurements and the number of preparations are included in the main body of the text and in the figure legends. Analyses were performed in Matlab and the statistical significance was assessed with Prism 8 (GraphPad Software, San Diego, CA, USA). Plots were generated in the Matlab and Prism softwares. Differences were analyzed with Student’s paired/unpaired t test or two-way ANOVA when appropriate and were considered significant at p<0.05. Details of the statistical tests used in each case are given in the text. Data expressed as mean ± s.e.m or s.d. as indicated.

## Acknowledgements

This work was supported by grants from the Swedish Research Council (2018-02692 and 2017-06092), the Torsten Söderberg foundation, AFA Försäkrings AB (170069), and the County Council of Östergötland (all to A.F.), the Fundación María Cristina Masaveu Peterson (to J.C. and R.C.) as well as the US National Institutes of Health (R01DC009645, to M.T.). We thank Anna Montell Magnusson for constructive criticism on earlier version of the manuscript.

## Author contributions

A.F. and S.P. conceived and designed the study. S.P. performed experiments, data acquisition and analysis of experimental data. A.F. and S.P. contributed to the experimental methodology. S.P. and A.F. wrote the manuscript, together with J.C. and B.V. B.V. assisted in collaboration and G.B., M.C., R.G.-A., A.Fo., C.D.-P., M.D., J.C., R.C., A.S. and M.T. performed genetic and/or clinical analyses of the patients. All authors commented on the manuscript.

## References

1. Kitajiri S, Fukumoto K, Hata M, Sasaki H, Katsuno T, Nakagawa T, Ito J, Tsukita S, Tsukita S. Radixin deficiency causes deafness associated with progressive degeneration of cochlear stereocilia. J Cell Biol. 2004;166(4):559–70.

2. Khan SY, Ahmed ZM, Shabbir MI, Kitajiri S, Kalsoom S, Tasneem S, Shayiq S, Ramesh A, Srisailpathy S, Khan SN, Smith RJ, Riazuddin S, Friedman TB, Riazuddin S. Mutations of the *RDX* gene gause nonsyndromic hearing loss at the DFNB24 locus. Hum Mut. 2007;28, 417–23.

3. Shearer AE, Hildebrand MS, Bromhead CJ, Kahrizi K, Webster JA, Azadeh B, Kimberling WJ, Anousheh A, Nazeri A, Stephan D, Najmabadi H, Smith RJ, Bahlo M. A novel splice site mutation in the RDX gene causes DFNB24 hearing loss in an Iranian family. Am J Med Genet A. 2009;149A(3):555–8.

4. Pataky F, Pironkova R, Hudspeth AJ. Radixin is a constituent of stereocilia in hair cells. Proc Natl Acad Sci U S A 2004;101, 2601–2606.

5. Shin JB, Krey JF, Hassan A, Metlagel Z, Tauscher AN, Pagana JM, Sherman NE, Jeffery ED, Spinelli KJ, Zhao H, Wilmarth PA, Choi D, David LL, Auer M, Barr-Gillespie PG. Molecular architecture of the chick vestibular hair bundle. Nat Neurosci. 2013;16, 365–374.

6. Hirono M, Denis CS, Richardson GP, Gillespie PG. Hair Cells Require Phosphatidylinositol 4,5-Bisphosphate for Mechanical Transduction and Adaptation. Neuron 2004;44, 309–320.

7. Pelaseyed T, and Bretscher A. Regulation of actin-based apical structures on epithelial cells. J Cell Sci. 2018;131(20).

8. Kunda P, Pelling AE, Liu T, Baum B. Moesin Controls Cortical Rigidity, Cell Rounding, and Spindle Morphogenesis during Mitosis. Curr Biol. 2008;18, 91–101.

9. Brownell WE, Bader CR, Bertrand D, de Ribaupierre Y. Evoked mechanical responses of isolated cochlear outer hair cells. Science 1985;227, 194–6.

10. Martin P, Mehta AD, Hudspeth AJ. Negative hair-bundle stiffness betrays a mechanism for mechanical amplification by the hair cell. Proc Natl Acad Sci USA 2000;97: 12026–12031.

11. Kennedy HJ, Crawford AC, Fettiplace R. Force generation by mammalian hair bundles supports a role in cochlear amplification. Nature 2005;433:880–883.

12. Meaud J, Grosh K. Coupling active hair bundle mechanics, fast adaptation, and somatic motility in a cochlear model. Biophys J 2011;100:2576 – 85.

13. Berninger E. Characteristics of normal newborn transient-evoked otoacoustic emissions: Ear asymmetries and sex effects. Int J Audiol. 2007;46, 661–69.

14. Cabanillas R, Dineiro M, Cifuentes GA, Castillo D, Pruneda PC, Alvarez R, Sanchez-Duran N, Capin R, Plasencia A, Viejo-Diaz M, Garcia-Gonzalez N, Hernando I, Llorente JL, Reparaz-Andrade A, Torreira-Banzas C, Rosell J, Govea N, Gomez-Martinez JR, Nunez-Batalla F, Garrote JA, Mazon-Gutierrez A, Costales M, Isidoro-Garcia M, Garcia-Berrocal B, Ordonez GR, Cadinanos, J. Comprehensive genomic diagnosis of non-syndromic and syndromic hereditary hearing loss in Spanish patients. BMC Med Genom. 2018;11:58

15. Warren RL, Ramamoorthy S, Ciganovic N, Zhang Y, Wilson TM, Petrie T, Wang RK, Jacques SL, Reichenbach T, Nuttall AL, Fridberger A. Minimal basilar membrane motion in low-frequency hearing. Proc Natl Acad Sci U S A 2016;113, 4304–10.

16. Strimbu CE, Prasad S, Hakizimana P, Fridberger A. Control of hearing sensitivity by tectorial membrane calcium. Proc Natl Acad Sci U S A 2019;116, 5756–5764.

17. Zhao H, Williams DE, Shin J-B, Brügger B, Gillespie PG. Large membrane domains in hair bundles specify spatially constricted radixin activation. J Neurosci. 2012;32, 4600–4609.

18. Gagnon LH, Longo-Guess CM, Berryman M, Shin J-B, Saylor KW, Yu H, Gillespie PG, Johnson KR. The chorlide intracellular channel protein CLIC5 is expressed at high levels in hair cell stereocilia and is essential for normal inner ear function. J Neurosci. 2006;26, 10188–10198.

19. Jacob S, Tomo I, Fridberger A, Boutet de Monvel J, Ulfendahl M. Rapid confocal imaging for measuring the sound-induced motion of the hearing organ. J Biomed Opt. 2007;12, 0211005.

20. von Tiedemann M, Fridberger A, Ulfendahl M, Boutet de Monvel J. Brightness-compensated 3-D optical flow algorithm for monitoring cochlear motion patterns. J Biomed Opt. 2010;15, 056012.

21. Kahsai AW, Zhu S, Wardrop DJ, Lane WS, Fenteany G. Quinocarmycin analog DX-52-1 inhibits cell migration and targets radixin, disrupting interactions of radixin with actin and CD44. Chem Biol. 2006;13, 973–83.

22. Kahsai AW, Zhu S, Fenteany G. G protein-coupled receptor kinase 2 activates radixin, regulating membrane protrusion and motility in epithelial cells. Biochim Biophs Acta 2010;1803, 300–310.

23. Jacob S, Pienkowski M, Fridberger A. The endocochlear potential alters cochlear micromechanics. Biophys J. 2011;100, 2586–94.

24. Oghalai JS, Tran TD, Raphael RM, Nakagawa T, Brownell WE. Transverse and lateral mobility in outer hair cell lateral wall membranes. Hear Res. 1999;135, 19–28.

25. Huang P, Zhang YH, Zheng XW, Liu YJ, Zhang H, Fang L, Zhang YW, Yang C, Islam K, Wang C, Naranmandura H. Phenylarsine oxide (PAO) induces apoptosis in HepG2 cells via ROS-mediated mitochondria and ER-stress dependent signaling pathways. Metallomics 2017;9, 1756–1764.

26. Bahloul A, Simmler MC, Michel V, Leibovici M, Perfettini I, Roux I, Weil D, Nouaille S, Zuo J, Zadro C, Licastro D, Gasparini P, Avan P, Hardelin JP, Petit C. Vezatin, an integral membrane protein of adherens junctions, is required for the sound resilience of cochlear hair cells. EMBO Mol Med. 2009;1, 125–38.

27. Fehon RG, McClachey AI, Bretscher A. Organizing the cell cortex: the role of ERM proteins. Nat Rev Mol Cell Biol. 2010;11, 276–287.

28. Tobin M, Chaiyasitdhi A, Michel V, Michalski N, Martin P. Stiffness and tension gradients of the hair cell’s tip-link complex in the mammalian cochlea. eLife 2019;8, 43473.

29. Chen F, Zha D, Fridberger A, Zheng J, Choudhury N, Jacques SL, Wang RK, Shi X, Nuttall AL. A differentially amplified motion in the ear for near-threshjold sound detection. Nat Neurosci. 2011;14, 770–74.

30. Ramamoorthy S, Zha D, Chen F, Jacques SL, Wang R, Choudhury N, Nuttall AL, Fridberger A. Filtering of acoustic signals within the hearing organ. J Neurosci. 2014;34, 9051–58.

31. Weichbold V, Nekahm-Heis D, Welzl-Mueller K. Universal newborn hearing screening and postnatal hearing loss. Pediatrics 2006;117, 631–6.

32. Young NM, Reilly BK, Burke L. Limitations of universal newborn hearing screening in early identification of pediatric cochlear implant candidates. Arch Otolaryngol Head Neck Surg. 2011;137, 230–4.

33. Dedhia K, Kitsko D, Sabo D, Chi DH. Children with sensorineural hearing loss after passing the newborn hearing screen. JAMA Otolaryngol Head Neck Surg. 2013; 139, 119–23.

34. Niparko JK, Tobey EA, Thal DJ, Eisenberg LS, Wang NY, Quittner AL, Fink NE; CDaCI Investigative Team. Spoken language development in children following cochlear implantation. JAMA 2010;303, 1498–506.

35. Boutet de Monvel J, Brownell WE, Ulfendahl M. Lateral Diffusion Anisotropy and Membrane Lipid/Skeleton Interaction in Outer Hair Cells. Biophys J. 2006;91, 364–381.

36. Bademci G, Foster J 2nd, Mahdieh N, Bonyadi M, Duman D, Cengiz FB, Menendez I, Diaz-Horta O, Shirkavand A, Zeinali S, Subasioglu A, Tokgoz-Yilmaz S, Huesca-Hernandez F, de la Luz Arenas-Sordo M, Dominguez-Aburto J, Hernandez-Zamora E, Montenegro P, Paredes R, Moreta G, Vinueza R, Villegas F, Mendoza-Benitez S, Guo S, Bozan N, Tos T, Incesulu A, Sennaroglu G, Blanton SH, Ozturkmen-Akay H, Yildirim-Baylan M, Tekin M. Comprehensive analysis via exome sequencing uncovers genetic etiology in autosomal recessive nonsyndromic deafness in a large multiethnic cohort. Genet Med. 2016;18, 364–71.

